# Disrupting myeloid-specific LXRα phosphorylation promotes FoxM1 expression and modulates atherosclerosis by inducing macrophage proliferation

**DOI:** 10.1101/228783

**Authors:** M. C Gage, N Bécares-Salles, R Louie, K Waddington, Yu Zhang, Thaís Tittanegro, S Rodriguez-Lorenzo, A Jathanna, B Pourcet, O Pello, J V De la Rosa, A Castrillo, I Pineda-Torra

## Abstract

Macrophages are key immune cells for the initiation and development of atherosclerotic lesions. However, the macrophage regulatory nodes that determine how lesions progress in response to dietary challenges are not fully understood. Liver X receptors (LXRs) are sterol-regulated transcription factors which play a central role in atherosclerosis by integrating cholesterol homeostasis and immunity. LXR pharmacological activation elicits a robust anti-atherosclerotic transcriptional program in macrophages that can be affected by LXRα S196 phosphorylation *in vitro*. To investigate the impact of these transcriptional changes in atherosclerosis development, we have generated mice carrying a Ser-to-Ala mutation in myeloid cells in the LDLR-deficient atherosclerotic background (M-S196A^Ldlr-KO^). M-S196A^Ldlr-KO^ mice fed a high fat diet exhibit increased atherosclerotic plaque burden and lesions with smaller necrotic cores and thinner fibrous caps. These diet-induced phenotypic changes are consistent with a reprogramed macrophage transcriptome promoted by LXRα-S196A during atherosclerosis development. Remarkably, expression of several proliferation-promoting factors including the proto-oncogene FoxM1 and its targets are induced by LXRα-S196A. This is consistent with increased proliferation of plaque-resident cells in M-S196A^Ldlr-KO^ mice. Moreover, disrupted LXRα phosphorylation increases expression of phagocytic molecules resulting in increased apoptotic cell removal by macrophages, explaining the reduced necrotic cores. Finally, the macrophage transcriptome promoted by LXRα-S196A under dietary perturbation is markedly distinct from that revealed by LXR ligand activation, highlighting the singularity of this post-translational modification. Overall, our findings demonstrate that LXRα phosphorylation at S196 is an important determinant of atherosclerotic plaque development through selective changes in gene transcription that affect multiple pathways.

## INTRODUCTION

Atherosclerosis is a chronic inflammatory process and the major pathology responsible for cardiovascular disease, which is now the leading cause of global mortality(1). This pathology results from the accumulation of lipids, immune cells and extracellular matrix within arterial walls, causing flow limitation(2). Atherosclerotic lesions progress and may rupture and thrombose, occluding the vessel and leading to myocardial infarcts or strokes. Macrophages are immune cells involved in most key pathways for the development of atherosclerosis including uptake of oxidized LDL, cholesterol efflux, foam cell and fatty streak formation, local proliferation, apoptosis, programmed removal of dead cells or efferocytosis, necrotic core formation and contribution to plaque stability(2).

Liver X receptors (LXRs) are ligand activated transcription factors that play vital roles in cholesterol homeostasis(3) and inflammation(4). LXRs are expressed as two isotypes; LXRα and LXRβ, which display 78% sequence homology, yet vary in their tissue expression and regulation(5). Both LXRs are endogenously activated by oxidized metabolites of cholesterol(6) and intermediates of the cholesterol biosynthesis pathway(7), as well as by various synthetic ligands(8). Pharmacological activation of these receptors has been demonstrated to modulate a range of lipid and inflammatory disorders(9). With regards to atherosclerosis, activating LXRs attenuates atherosclerosis progression(3) via promotion of cholesterol efflux through lipid-laden macrophages present in the atherosclerotic lesions, inhibition of vascular inflammation(4) and possibly by affecting other aspects of lipid metabolism(10). Additionally, ligand-activated LXR promotes CCR7-dependent plaque regression(11). Functional studies in macrophages further indicate that LXRα is required for a robust anti-atherosclerotic response to LXR ligands and LXRα plays a selective role in limiting atherosclerosis in response to hyperlipidemia(12).

LXRα transcriptional activity can be modulated by several posttranslational modifications(5) including phosphorylation at serine (S) 198 in the human sequence, corresponding to S196 in the mouse orthologue. We have demonstrated that modulation of LXRα phosphorylation significantly modifies its target gene repertoire in macrophage cell lines overexpressing the receptor, thereby altering pathways known to be relevant to the development of atherosclerosis(13, 14). Interestingly, we previously showed phosphorylated S196-LXRα is present in progressive atherosclerotic lesions(14) suggesting LXRα phosphorylation at this residue could be important for the development of atherosclerotic plaques. However, the specific contribution of myeloid LXRα phosphorylation to atherosclerosis development remains unknown.

To investigate this, we have generated a mouse model specifically expressing a Ser-to-Ala phosphorylation mutant of LXRα in myeloid cells (M-S196A) in the LDLR-deficient (Ldlr-KO) atherosclerotic background (M-S196A^Ldlr-KO^). Disrupting LXRα phosphorylation in myeloid cells including macrophages promotes plaque burden, yet modulates plaque phenotype to acquire distinctive characteristics such as smaller necrotic cores and thinner fibrous caps encapsulating the lesions. These phenotypic changes are consistent with a reprogrammed macrophage transcriptome. Notably, cell cycle progression and proliferation pathways are markedly induced in M-S196A^Ldlr-KO^ macrophages, specifically the expression of the FoxM1 transcription factor and several of its targets. This is associated with increased lesion-resident cell proliferation in the LXRα phospho-mutant mice. In addition, changes in the expression of various phagocytic molecules result in enhanced macrophage efferocytosis thus explaining the reduced necrotic cores present in M-S196A mice. Interestingly, most of the phosphorylation sensitive genes identified are not subject to LXR ligand regulation and we show that global transcriptional changes in response to impaired LXRα phosphorylation under dietary perturbation are markedly distinct from those revealed by ligand activation. Overall, these findings demonstrate LXRα phosphorylation at S196 determines atherosclerotic plaque progression by promoting changes in local cell proliferation, efferocytosis and necrotic core formation.

## RESULTS

### Impaired myeloid LXRα phosphorylation promotes atherosclerosis

To investigate the impact of macrophage LXRα phosphorylation on the development of atherosclerosis we generated a new mouse model expressing a serine to alanine mutation at residue 196 in LXRα in myeloid cells (M-S196A) on a pro-atherosclerotic (LDLR-deficient or Ldlr-KO) background (M-S196A^Ldlr-KO^) (Fig. S1A). Effective expression of Cre-driven targeting construct introducing S196A knock-in in the sense strand was demonstrated in M-S196A^Ldlr-KO^ compared to WT^Ldrl-KO^ control littermates (Fig. S1B). Mice were fed a fat-rich Western diet (WD) to accelerate plaque progression. M-S196A^Ldlr-KO^ mice developed normally and no change in body weight before, during or after Western diet (WD) feeding was observed (Fig. S1C-E). There were no detectable changes in basal metabolic characteristics including total cholesterol, HDL or LDL/VLDL levels and amount of triglycerides, free fatty acids and insulin in the plasma of M-S196A^Ldlr-KO^ compared to WT^Ldlr-KO^ (Fig. S2A-G). Interestingly, M-S196A^Ldlr-KO^ mice showed a significant increase in atherosclerosis plaque burden in their aortas as measured by *en face* oil red O staining (Fig. 1A) and aortic root plaque coverage (Fig. 1B). This was however, not associated with changes in the levels of CD68+ positive cells in the lesions (Fig. 1C).

**Figure 1.**
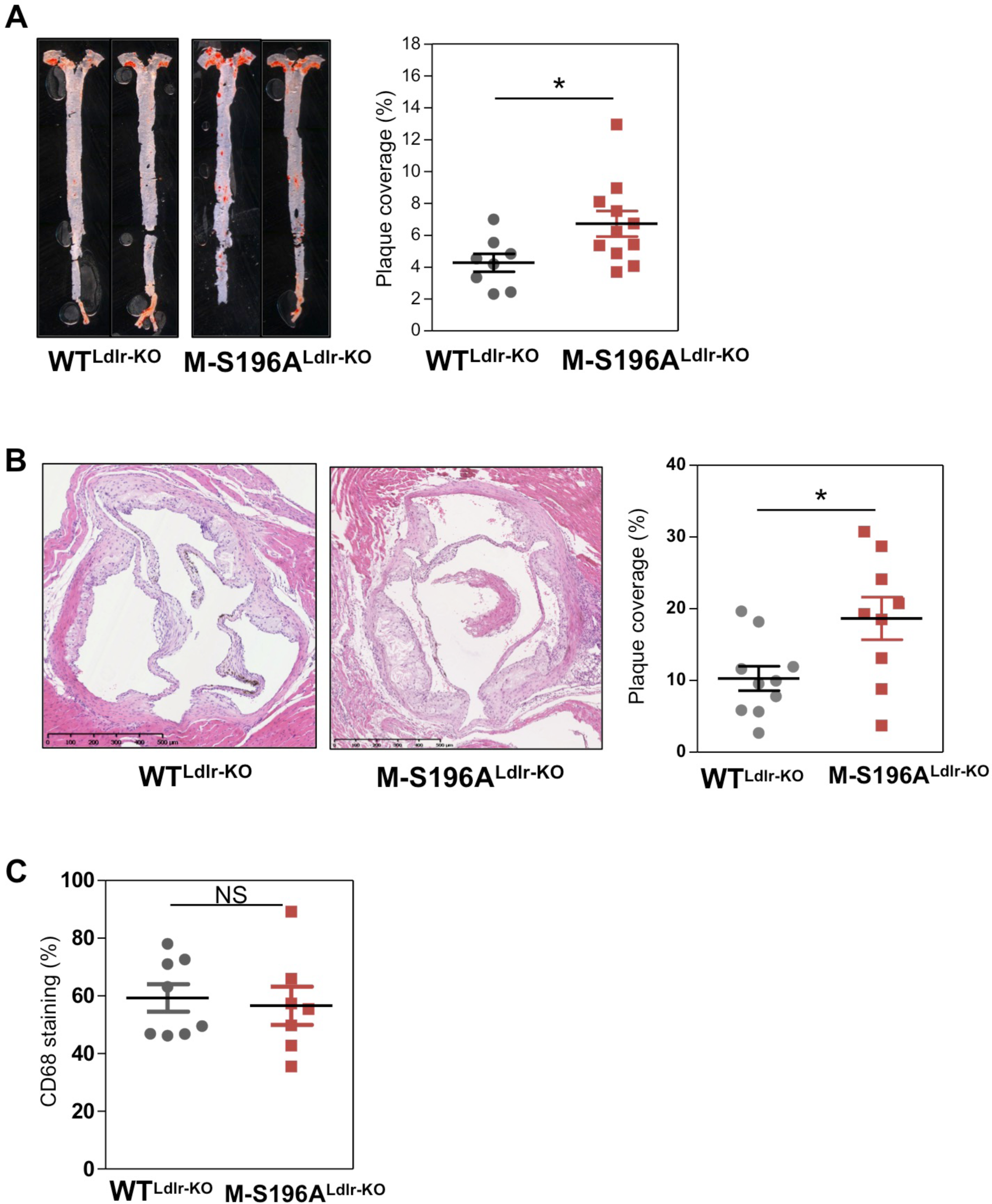
M-S196A^Ldlr-KO^ mice develop increased atherosclerosis on a Western diet. (**A**) (*Left*) En face Oil Red O-stained whole aortas (n=8-11/group), representative images shown. (*Right*) Quantification of stained areas as *%* plaque coverage for each genotype. (**B**) (*Left*) Haemotoxylin and Eosin (H&E)-stained aortic roots (n=9-10/group), scale bar at 500 μm representative images shown. (*Right*) Quantification of stained areas as % plaque coverage for each genotype. (**C**) CD68 staining of aortic roots (n=7-8/group). Data are means ± SEM. *p<0.05, relative to WT^Ldlr-KO^.

### Changes in LXRα phosphorylation at Ser196 reprogram global macrophage gene expression in the context of diet-induced atherosclerosis

To explore in more detail the pathways underlying the changes observed in atherosclerosis development we investigated the transcriptomic profiles of macrophages differentiated from the bone marrows of mice exposed to the WD. RNA-seq analysis revealed significant genome wide changes in transcript levels (Fig. 2A, B). LXRα-S196A significantly induced (460) or reduced (210) gene expression. Hallmark pathway analysis identified G2M checkpoint and E2F targets to be markedly enriched indicating cell cycle and cell proliferation pathways are induced in the mutant-expressing macrophages (Fig 2C and Fig. S3A,B). This was further confirmed by Reactome pathway analysis (Table S1). Several genes involved in these processes were regulated over 2-fold including cell proliferation marker *Mik67* or Ki67 (4.58-fold, p=3.46E-43) (Fig 2D). Concomitant to these changes in cell proliferation genes, there was a substantial reduction in the expression of genes associated with the immune response (Fig 2C,E, Fig. S3C and Table S2). We observed opposing changes in the expression of chemokine receptors involved in monocyte trafficking to atherosclerotic lesions and some of the chemokines they bind to(15) (Fig. 2F). For example, expression of the chemokine receptors *Ccr1* (1.43-fold, P=0.004), *Ccr2* (1.73-fold, P=3.8×10^-15^) and *Cx3cr1* (1.8-fold, P=0.0008) was increased whereas Ccr5 (0.55-fold, P=6.19×10^-17^) expression was diminished in LXRα-S196A macrophages compared to WT macrophages. Such differential expression of chemokine receptors and their ligands may explain the lack of change in the overall number of CD68+ cells retained in the plaques of M-S196A^Ldlr-KO^ mice.

**Figure 2.**
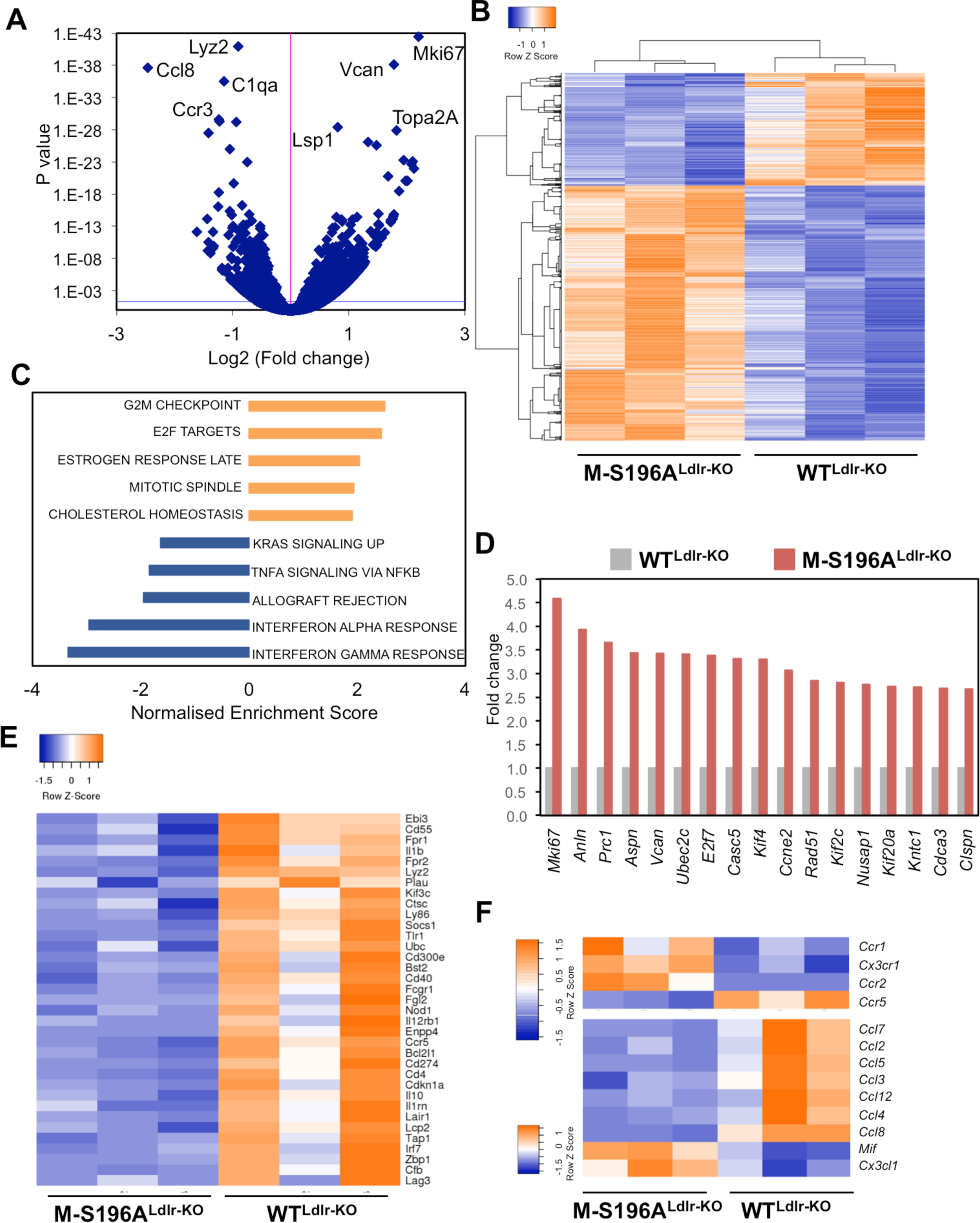
Changes in LXRα phosphorylation reprogram macrophage gene expression. (**A**) Volcano plot of log_2_ ratio vs p-value of differentially expressed genes comparing 12 week WD-fed M-S196A^Ldlr-KO^ to WT^Ldlr-KO^ BMDM (n=3/group). Blue line indicates adjusted p-value threshold of 0.04 (Wald Test for logistic regression. (**B**) Clustered heatmap of RNAseq gene counts in WD-fed macrophages (n=3 mice/group). (**C**) GSEA analysis showing enriched pathways in M-S196A^Ldlr-KO^ macrophages derived from HALLMARK gene sets. (**D**) Fold-change of RNAseq gene counts in M-S196A^Ldlr-KO^ compared to WT^Ldlr-KO^ (set as 1) (n=3/genotype) for top induced genes (≥2-fold expression, p ≤0.01) involved in cell proliferation. **E**) Heatmap of RNAseq gene counts of immune response genes downregulated by S196A in WD-fed macrophages (n=3 mice/group). **F**) Heatmap of RNAseq gene counts of (*Top*) chemokine receptor and (*Bottom*) chemokine ligand genes showing differentially expressed genes in S196A WD-fed macrophages (n=3 mice/group). For all heatmaps blue and orange depicts upregulated and downregulated genes respectively and only genes showing ≥1.3-fold change with p ≤0.01 are shown.

### M-S196A induces expression of FoxM1 and lesion-resident cell proliferation

Examination of the RNA-seq datasets revealed LXRα-S196A cells expressed almost 3-fold (p=7.76E-14) more proto-oncogene FoxM1 compared to macrophages expressing wild type LXRα (Fig. 3A,B). This was also the case for several FoxM1 target genes(16) (Fig. 3A,B). While LXRα activation was previously shown to inhibit cell proliferation via inhibition of FoxM1 in hepatic carcinoma cells(17), its regulation in macrophages has never been documented. LXRs modulate gene transcription by heterodimerising with the Retinoid X Receptor (RXR) and binding to specific DNA sequences termed LXR response elements (LXREs) in the transcriptional regulatory regions of their target genes(18). Notably, specific LXRα occupancy was observed at the *FoxM1* gene in macrophages further indicating that *FoxM1* is an LXRα target in these cells (Fig. S4A). The enhanced levels of several pro-mitotic genes suggested cell proliferation could be altered in M-S196A^Ldlr-KO^ macrophages. Indeed macrophages expressing the LXRα-S196A mutant showed about 20% increase in proliferation in culture measured as Ki67 levels (Fig. S4B). Recent studies have highlighted the important role local macrophage proliferation plays in lesion development(19). Consistent with a significant increase in the regulation of *FoxM1* and other genes involved in cell cycle pathways, increased proliferation of lesion-resident cells as measured by Ki67 staining was observed in the atherosclerotic plaques of M-S196A^Ldlr-KO^ mice (Fig. 3C,D) which was associated with increased nuclei content (Fig. S5A) compared to WT^Ldlr-KO^. This strongly suggests that enhanced local proliferation in the plaques could contribute to increased plaque size exhibited by M-S196A^Ldlr-KO^ mice as has been postulated(20-22).

**Figure 3.**
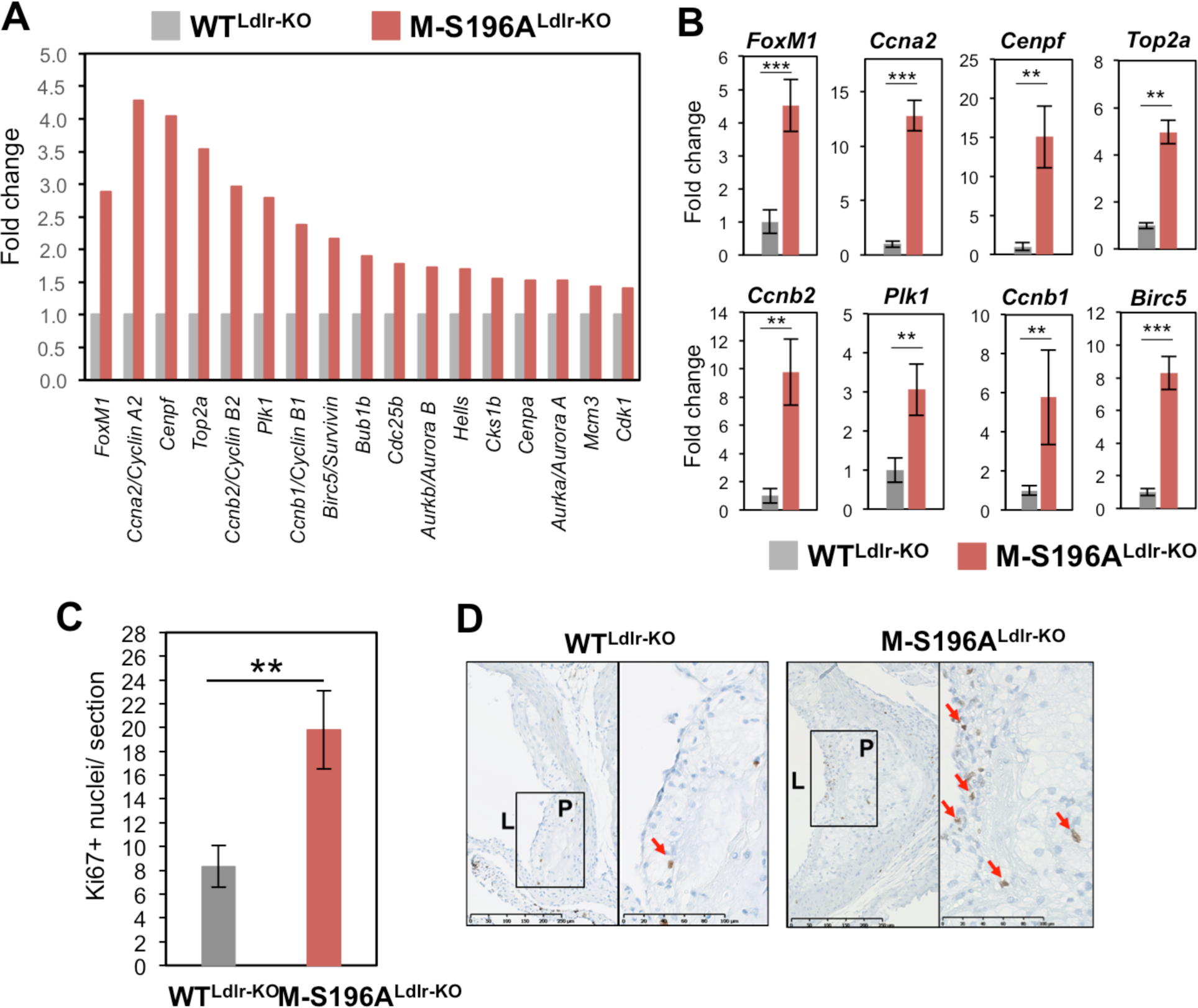
Impaired macrophage LXRα phosphorylation induces FoxM1 expression and increases plaque cell proliferation. (**A**) Fold-change of RNAseq gene counts in WD-fed M-S196A^Ldlr-KO^ compared to WD-fed WT^Ldlr-KO^ macrophages (set as 1) for FoxM1 and FoxM1 target genes with ≥1.3-fold expression, p ≤0.01, n=3/genotype. (**B**) RT-qPCR analysis of FoxM1 and top FoxM1 regulated targets in WD-fed WT^Ldlr-KO^ and M-S196A^Ldlr-KO^ macrophages. Normalized data shown relative to WT^Ldlr-KO^ (set as 1) as mean ± SEM, n=3, **p<0.01 or ***p<0.001. (**C**) Quantification of Ki67 positive nuclei in WD-fed WT^Ldlr-KO^ and M-S196A^Ldlr-KO^ plaques (n= 6-10 mice/group). (**D**) Representative images of plaques exhibiting Ki67 positive nuclei, scale bar at 250 μm.

### M-196A^Ldlr-KO^ mice display phenotypic changes in their necrotic cores and fibrous caps

The observed changes in gene expression suggest a complex interaction of pathways involved in the progression of atherosclerosis. Unexpectedly, despite their larger size, size-matched atherosclerotic lesions in M-S196A^Ldlr-KO^ mice display smaller necrotic cores (Fig. 4A). Programmed cell removal or efferocytosis has been shown to strongly impact the formation of necrotic cores in advanced plaques(23). In agreement with this, macrophage engulfment of apoptotic cells was significantly increased (Fig. 4B and Fig. S5B). Further interrogation of the LXRαS196A-regulated transcriptome showed differential expression of several pro- and antiphagocytic molecules (Fig. 4C). This includes *Ccr2* (1.73-fold, P=3.8×10^-15^), *Gpr132* (1.64-fold, P=1.73×10^-10^), Itgb3 (1.59-fold, P=0.007) and *Mfge8* (1.37-fold, P=7.46×10^-5^) known to promote efferocytosis(23) as well as molecules known to render apoptotic cells resistant to efferocytosis such as *Cd47(24)* (0.75-fold, P=0.0001) and *Tnf* (0.71-fold, P=0.0004) in M-S196A^Ldlr-KO^ macrophages. This is consistent with the enhanced efferocytosis observed in these cells. Another important morphological feature of atherosclerotic lesions influenced by macrophages is the thinning of the protective collagenous scar surrounding it or fibrous cap(25). Interestingly, M-S196A^Ldlr-KO^ lesions show reduced fibrous cap thickness with overall smaller fibrous cap areas (Fig. S5C,D). This could result from the diminished expression of several collagen species including *Col1a1* (0.6-fold, P=9.3×10^-04^), *Col1a2* (0.7-fold, P=9.9×10^-03^), *Col3a1* (0.6-fold, P=4.0x10^-03^), *Col5a1* (0.6-fold, P=3.4×10^-03^), and *Col6a1* (0.5-fold, P=4.2×10^-05^) and increased levels of matrix degrading molecules such as *Mmp8* (1.57-fold, P=6.1×10^-10^) and *Mmp12* (1.48-fold, P=8.8×10^-6^). Overall, this data highlights the complex phenotypic changes present in atherosclerotic lesions resulting from changes in LXRα phosphorylation.

**Figure 4.**
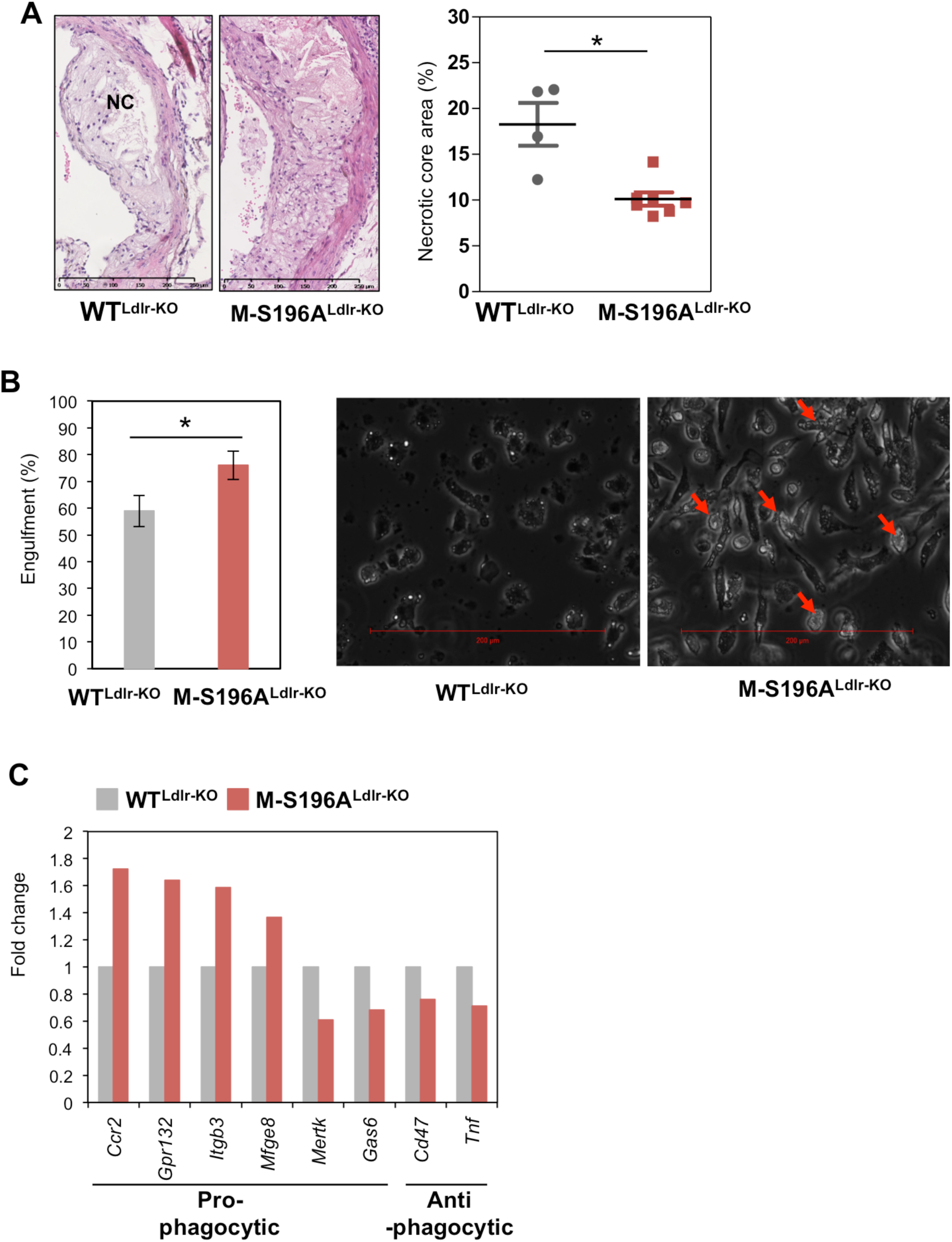
M-S196A^Ldlr-KO^ mice show decreased plaque necrotic cores and increased efferocytosis capacity. (**A**) (*Left*) H&E-stained mature plaques ‘NC’ depicts necrotic core (n= 4-6/group), representative images shown, scale bar at scale bar at 250 μm. (*Right*) Quantification of H&E-stained areas for each genotype. (**B**) (*Left*) Engulfment of apoptotic Jurkat cells (n=6/group), (*Right*) Representative images for each genotype shown, scale bar at 200 μm. (**C**) Fold-change of RNAseq gene counts in WD-fed M-S196A^Ldlr-KO^ compared to WD-fed WT^Ldlr-KO^ macrophages (set as 1) for pro- and anti-phagocytic genes, p **≤**0.01 n=3/genotype.

### Disrupted LXRα phosphorylation at Ser196 alters ligand responses in macrophages

Our findings indicate that in the context of an atherogenic diet, changes in LXRα phosphorylation modulate the macrophage transcriptome and promote atherosclerotic plaque burden. To further understand the magnitude of the transcriptional changes imposed by the LXRα phosphorylation mutant, we next examined whether WT and S196A expressing macrophages respond differently to an LXR ligand and explored the differences in global transcript changes between ligand activation and reduced LXRα phosphorylation. RNA-seq analysis was performed on bone marrow-derived macrophages from M-196A^Ldlr-KO^ mice exposed to WD for 12 weeks and cultured in the presence of vehicle or LXR ligand GW3965. GW ligand activation promoted changes in macrophage gene expression that were different in cells expressing the S196A mutant compared to WT macrophages (Fig. S6A-C). GSEA analysis revealed the pathways subject to changes in LXRα phosphorylation in the presence of the LXR ligand are similar to those seen in the absence of GW (Fig. S6D). For instance, genes involved in nuclear division and cell cycle remained strongly induced by LXRα-S196A, further emphasizing the importance of LXRα phosphorylation in the modulation of these pathways. Remarkably, it became apparent that while a small subset of genes were differentially regulated by the mutant only in the context of the ligand (94 induced and 50 reduced compared to WT cells), most differences in gene expression were observed in the absence of ligand (Fig. 6A,B). Additionally, our datasets also showed that ligand responses were similar in both WT and mutant expressing macrophages (Fig. 6C,D). However, the identity of the genes regulated was distinct, with only 47 genes (about half of the total number) being regulated by both WT and S196A forms of LXRα (Fig. 6C). Further analysis revealed that both the magnitude of the response and the identity of the genes were strikingly different between the response to the ligand (regulation by GW in either WT or S196A cells) and to phosphorylation (modulation by LXRα-S196A compared to WT) (Fig. 6E,F). This highlights the significance of phosphorylation in rewiring the LXR-modulated transcriptome.

Finally, we investigated whether this dichotomy between ligand and phosphorylation-induced responses was apparent in the regulation of the phosphorylation-sensitive gene *FoxM1* and some of its target genes. *FoxM1* was not significantly affected by exposure to the LXR ligand in WT^LdlrKO^ cells (Fig. 5G). By contrast, GW3965 activation markedly reduced *FoxM1* mRNA levels in M-S196A^Ldlr-KO^ macrophages (Fig. 5G). This regulatory pattern was recapitulated by most *FoxM1* targets examined (Fig. 5G and Fig. S6E). In addition, established transcriptional regulators of *FoxM1* were strongly *(Top2a, Rad51, Check2)* or moderately *(Melk,)* induced in M-S196A^Ldlr-KO^ cells in unstimulated conditions compared to WT^LdlrKO^ cells (Fig. 5H). Consistent with the known anti-proliferative effects of LXR ligands, the expression of these genes was strongly attenuated by GW3965, particularly in the mutant cells. Other phosphorylation-sensitive genes implicated in cell cycle progression mimic this mode of regulation (Fig S6F).

**Figure 5.**
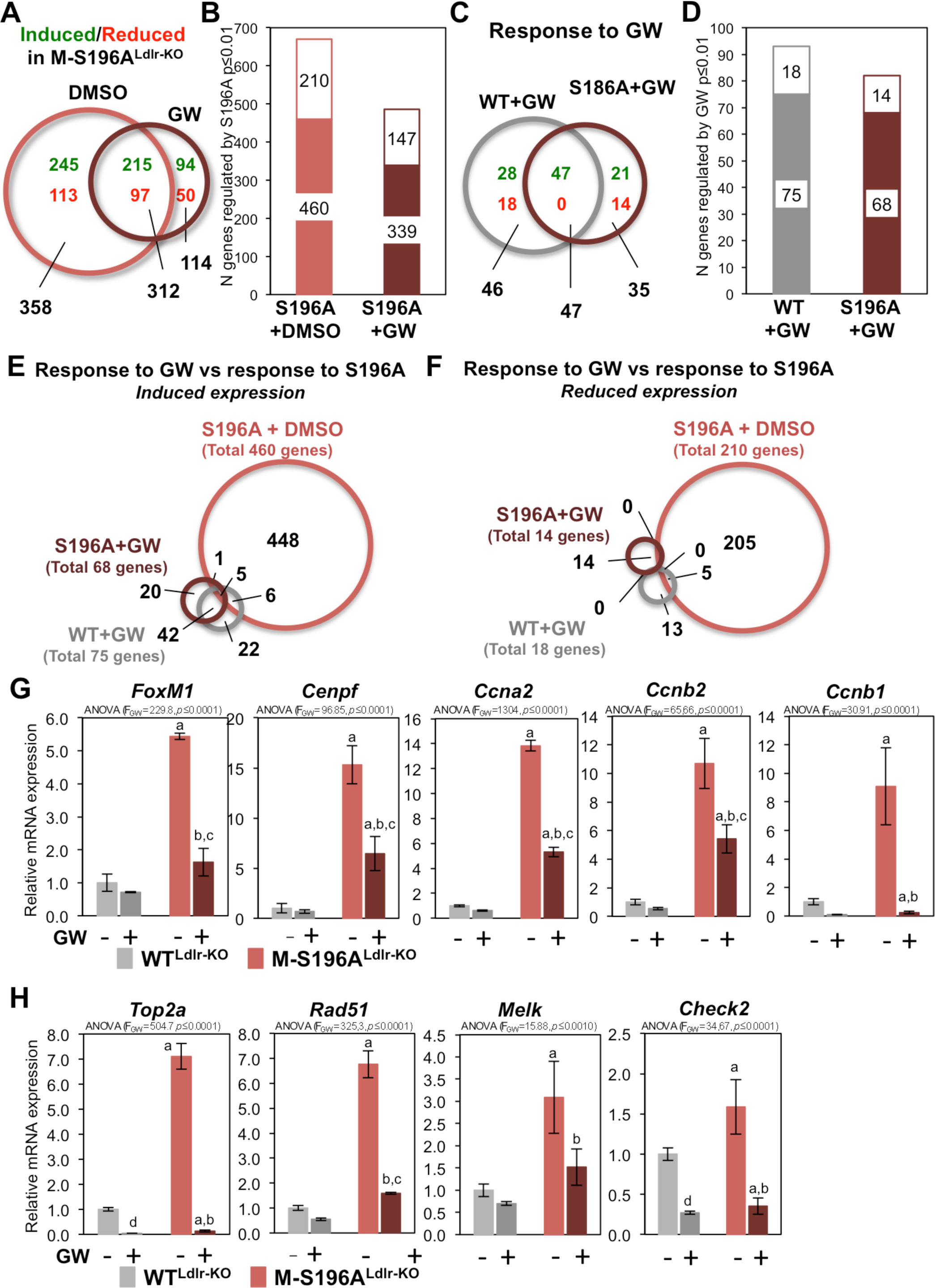
Macrophage transcriptional reprogramming in response to changes in LXRα phosphorylation is fundamentally different from ligand activation responses. (**A**) M-S196A^Ldlr-KO^ and WT^Ldlr-KO^ macrophages from WD-fed mice exposed to 1μM GW3965 (GW) (n=3/group). Venn diagram of genes regulated by LXRαS196A compared to LXRαWT. Numbers of genes showing induced or reduced expression are depicted in green and red, respectively. (**B**) Number of genes showing ≥1.3-fold regulation changes (p ≤0.01) due to LXRα-S196A expression. Solid and open bars represent genes that exhibit induced or reduced expression respectively. (**C**) Venn diagram of genes regulated in response to GW treatment (DMSO compared to GW) for each genotype. Numbers of genes exhibiting induced or reduced expression shown as in panel C. (**D**) Number of genes showing ≥1.3-fold regulation changes (p ≤0.01) upon GW treatment. Bars show gene numbers as in panel D. (**E**) Venn diagram showing number of genes induced by LXRα-S196A or LXRα-WT in GW-treated macrophages compared to DMSO-treated cells (showing ligand responses) and genes induced by LXRα-S196A compared to LXRα-WT cells under basal conditions (S196A+DMSO-showing responses to impaired LXRα phosphorylation). (**F**) Venn diagram as in panel G comparing number of genes reduced by GW ligand or by impaired LXRα phosphorylation. (**G**) RT-qPCR analysis of *FoxM1* and its target genes in WD-fed WT^Ldlr-KO^ and M-S196A^Ldlr-KO^ macrophages. Normalized data shown relative to WT^Ldlr-KO^ as mean ± SD, n=3, (a) p≤0.001 compared to WT-DMSO, (b) p≤0.001 compared to S196A-DMSO, (c) p≤0.001 compared to WT-GW. (**H**) mRNA expression of *FoxM1* regulators. Data and statistical analysis as in G.

Overall, our findings suggest that LXRα phosphorylation at Ser196 is a powerful means of regulating LXRα transcriptional activity that has important consequences for macrophage biology and for the progression of a metabolic, inflammatory and proliferative disease such as atherosclerosis.

## DISCUSSION

The macrophage regulatory nodes that determine how the atherosclerotic lesion progresses in response to dietary challenges are not fully understood. Liver X receptors have key roles in the regulation of macrophage lipid homeostasis and inflammation and as such they strongly modulate the progression of metabolic diseases such as atherosclerosis(3). The importance of these receptors in disease development has been mainly gleaned from studies evaluating the consequences of its pharmacological or genetic manipulation. However, it remained unknown whether alternative modulation of the activity of these receptors, for instance, by altering post-translational modifications of the receptor, could shape the pro-atherogenic responses of fat-rich diets thus altering disease development. We previously showed LXRα is phosphorylated in cholesterol-loaded macrophages and in progressive atherosclerotic plaques(13). We have now explored the impact of LXRα phosphorylation on atherosclerosis development by expressing an LXRα Ser-to-Ala mutant, previously shown to disrupt LXRα phosphorylation(13), specifically in myeloid cells on the LDLR null background (M-LXRαS196A^Ldlr-KO^).

LXRαS196A expression in cells of the myeloid lineage, including macrophages, increases atherosclerotic plaque burden (Fig. 1). This is consistent with an enhanced number of proliferating lesion-resident cells (Fig. 3C,D) in M-LXRαS196A^Ldlr-KO^ mice and the up-regulation of genes driving cell cycle progression in macrophages, particularly at the G2/M checkpoint, accounting for up to 15% of the total changes in gene expression exerted by the phosphorylation mutant (Fig. 2C). During the past decade established paradigms of atherosclerotic plaque formation and progression have been revisited and local proliferation of macrophages has been demonstrated to be an important driver of atherosclerosis development in advanced atherosclerotic plaques(20, 22, 26). However, the specific players modulating macrophage proliferation in the context of atherosclerosis remain poorly understood. Proliferation of lesional macrophages has been linked to up-regulation of the scavenger receptor Msr1(20, 27) but a defined mechanism has remained elusive. Our findings now indicate that modulation of LXRα phosphorylation plays an important role in this process.

LXRs are known modulators of cell proliferation in other disease conditions where cell proliferation is critical, including cancer(28). For instance, LXR activation inhibits proliferation of B and T cells and macrophages(29-31) and several cancer cell lines including prostate (LNCaP)(32), breast (MCF7)(33) and colon (HTC111)(34). Identified anti-proliferative mechanisms by classic LXR agonists appear to be independent of the lipogenic activity of LXR(33), but rather linked to βcatenin activity(34), cyclins(32) and sterol metabolism(31). In contrast to these inhibitory effects, LXR activation was recently reported to enhance proliferation of neural progenitor cells in a MEK-ERK pathway dependent manner(35). Inverse agonism of LXR with a novel synthetic agonist has also shown promise as a potential cancer treatment though inhibition of lipogenesis, glycolysis and by regulating the expression of key glycolytic and lipogenic genes(28). However this is the first time LXR is shown to target cell cycle promoting factors in an atherosclerotic context.

The underlying mechanisms explaining the reported LXR anti-proliferative actions may be cell-specific. The inhibitory effects of LXR ligands on M-CSF-stimulated macrophage proliferation involve the down-regulation of the cyclin D kinase regulators cyclin D1 (*Ccnd1*) and B1 (*Ccnb1*)(30). Both cyclin regulators are inhibited by LXRα through the transcriptional repression of FoxM1 in hepatic carcinoma cells(17). Conversely, LXR antagonism with a sulphated oxysterol promotes hepatic proliferation in part through the induction of FoxM1(36). FoxM1 is an essential proliferation-associated transcription factor found overexpressed in numerous solid tumors(37-40). Its expression is restricted to actively dividing cells, and is regulated in a cell-cycle dependent manner by a wide range of proliferative signals(41). FoxM1 levels are induced in the G1-phase, continuing throughout the S- phase and reaching maximum expression in the G2/M-phase(42-45). We now demonstrate that chronic disruption of LXRα phosphorylation in macrophages enhances FoxM1 expression and several of its associated regulators and targets driving cell cycle progression(16) (Fig. 3A,B). This is associated with increased lesional cell proliferation (Fig. 3C,D) and cell content (Fig. S5A), consistent with larger atherosclerotic lesions. Amongst FoxM1 regulators, Bub1b (or BubR1, increased 1.9 fold, P=1.26×10^-6^ in the LXRα-S196A-expressing macrophages) and has been shown to alter atherosclerosis, as impaired expression of BubR1 results in decreased macrophage proliferation and attenuated atherogenesis(47). Furthermore, LXRα-S196A macrophages show decreased mRNA expression of the FoxM1 inhibitor SASH1(46) (0.8-fold, P=1.33×10^-4^), which was implicated in the development of atherosclerosis in people that smoke through its monocytic up-regulation. Overall, our findings further highlight the importance of this set of molecules in atherosclerosis progression.

Importantly, we show evidence that reduced LXRα phosphorylation during atherosclerosis development reprograms the macrophage transcriptome (Fig. 2). Global gene expression analysis revealed that in these cells, most genes are sensitive to the expression of the LXRα phosphomutant in the absence of LXR ligand stimulation, with only an additional 114 genes being modulated by the mutant in the presence of the GW3965 ligand compared to the 670 LXRα phosphorylation sensitive genes under basal conditions. This suggests that modulation by LXRα-S196A expression is different from the regulation by ligand activation of the receptor. Indeed, comparison of the various datasets evidenced that the transcriptomic rewiring in response to impaired LXRα phosphorylation in the mutant cells is remarkably distinct and does not merely phenocopy ligand responses (Fig. 5E,F) thus highlighting the importance of this post-translational modification in modulating the activity of LXRα in the context of a metabolic and inflammatory disease. An example for this is the regulation of FoxM1 expression and its regulated pathways, which are induced in LXRα-S196A-expressing cells (Fig. 3A). Previous reports have linked LXRα activation with FoxM1 repression in hepatic carcinoma cells(17). We observed that FoxM1 inhibition by GW3965 is recapitulated in macrophages from mice exposed to a high fat diet that are developing atherosclerotic plaques (Fig. 5G). However, this is preferentially observed in the LXRα phosphorylation mutant-expressing cells. In fact, most FoxM1 targets and modulators as well as other factors involved in the G2/M cell cycle transition mirror this pattern of regulation (Fig. 5G,F and Fig. S6E,F). This suggests that in macrophages, the anti-proliferative effects of LXR ligands are enhanced when LXRα phosphorylation is disrupted. Intriguingly, these transcriptomic changes are observed in cells that have been differentiated and cultured *in vitro* from precursor cells exposed to the pro-atherosclerotic environment in the bone marrow of WD-fed mice. This suggests that cells retain the "memory” of this context in culture. Indeed, the concept of epigenetic memory of innate immune cells such as macrophages has been proposed to play an important role in modulating immune responses(48-51). Although this has been mainly studied in the context of pro-inflammatory stimuli and the regulation of pro-inflammatory gene expression, there are also reports showing a distinct metabolic environment (for instance in type 2 diabetes) can epigenetically imprint bone marrow progenitor cells that can derive into a "pre-programmed macrophage state associated with changes in gene expression(51). Outside the scope of this study, future investigations will help establish changes in the epigenome of LXRα-S196A bone marrow-derived macrophages and how they affect metabolic, proliferative and inflammatory pathways in the context of atherosclerosis.

Beyond changes in cell proliferation, the enhanced plaque burden in M-LXRαS196A^Ldlr-KO^ mice is likely the result of the complex modulation of additional pathways relevant for atherosclerosis development. LXRα promotes the expression of factors important for macrophage efferocytotic capacity such as the MerTK receptor for apoptotic cells(52) which can influence the formation of the necrotic core and plaque stability(53). Despite the overall increase in atherosclerosis, we found that size-matched advanced plaques from M-S196A^Ldlr-KO^ mice exhibited significantly reduced necrotic cores (Fig. 4A) consistent with the increased capacity of M-S196A^Ldlr-KO^ macrophages to engulf apoptotic cells (Fig. 4B& S5A). Transcriptomic profiling revealed that in addition to the reduced expression of Mertk and its ligand Gas6, several genes known to promote phagocytosis were significantly upregulated in M-S196A^Ldlr-KO^ cells, including the pro-phagocytotic receptors Ccr2(54), Gpr132(55), Itgb3(56) and the bridging molecule Mfge8(57) (Fig. 4C). This differential expression could explain the increased phagocytic ability of the LXRα phospho-mutant expressing macrophages (Fig. 4B).

In summary, we have shown that disrupting LXRα phosphorylation in cells of the myeloid lineage, affects the development of atherosclerosis that could be explained through altered cell proliferation and efferocytosis. We also show that chronically modulating LXRα phosphorylation reprograms gene regulation in macrophages under basal conditions and significantly affects their response to ligand stimulation. These findings add to our fundamental knowledge of how LXRα activity can be regulated and introduce novel functional consequences of its modification by phosphorylation which should be heeded to manipulate these receptors for the design of novel cardiovascular therapies.

## MATERIALS AND METHODS

### Mice

The S196A floxed (S196A^fl/fl^) mouse line was generated by Ozgene Pty Ltd (Bentley WA, Australia). The genomic sequence for the murine LXRα (Nr1h3) gene was obtained from the Ensembl Mouse Genome Server (https://www.ensembl.org/Mus_musculus/), Ensembl gene ID: ENSMUSG00000002108. The mutant fragment, located on Exon 5, contains a serine-to-alanine mutation at Ser196 introduced by site-directed mutagenesis. The point-mutant exon was delivered into an intronic site inside the targeting vector, placed in opposite orientation and thus without coding capacity (Fig. S1A). The targeting construct was electroporated into the Bruce4 C57BL/6 ES cell line. Homologous recombinant ES cell clones were identified by Southern hybridization and injected into BALB/cJ blastocysts. Male chimeric mice were obtained and crossed to C57BL/6J females to establish heterozygous germline offsprings on a pure C57BL/6 background. The germline mice were crossed to a FLP Recombinase mouse line(58) to remove the FRT flanked selectable marker cassette (Flp’d mice (Flp^+/+^). Flp^+/+^ mice are homozygous for FL allele containing LXRα WT exon 5 (Ex5) and LXRα S196A exon 5 of the LXRα gene in opposite orientation flanked by lox sites sensitive to Cre recombinase activity. These mice express LXRα WT but switch to LXRα S196A expression in the presence of CRE recombinase. Flp^+/+^ mice were crossed with (1) a C57BL/6 homozygous Ldl receptor null (Ldlr^-KO^) mouse strain, to generate a Flp^+/+^Ldlr^-KO^ strain and (2) a C57BL/6 homozygous LysMCre (LysMCre^+/+^) strain to generate a Flp^+/+^LysMCre^+/−^ strain. The two resulting lines were then crossed to generate Flp^+/+^Ldlr^-KO^LysMCre^+/−^. Cre recombinase expression under direction of the LysM promoter in Flp^+/+^Ldlr^-KO^LysMCre^+/−^ results in the switch to LXRα S196A expression in myeloid cells, these mice (M-S196A^Ldlr-KO^) were compared to littermate non- CRE expressing mice (Flp^+/+^Ldlr^-KO^LysMCre^-/-^ or WT^Ldlr-KO^) which were used as controls in our study. The Ldlr-KO and LysMCre strains were purchased from The Jackson Laboratory (stock numbers 002207 and 004781, respectively) Animals were housed together and maintained in a pathogen-free animal facility in a 12-h light-12h dark cycle. All procedures were carried under the UK’s Home Office Animals (Scientific Procedures) Act 1986.

### Genotyping

Mice were genotyped by PCR analysis of ear biopsies using Jumpstart Taq DNA Polymerase (Sigma Aldrich) and the following primers: For S198A knock-in, primers R2 and WT identify the wild-type allele (642bp) and FL allele (656bp), primers R2 and SA identifly the mutat allele of LXRα S196A knock-in mice (656 bp): SA 5’ GGTGTCCCCAAGGGTGTCCG wild-type 5’ GGTGTCCCCAAGGGTGTCCT, R2 5’ AAGCATGACCTGCACACAAG, Ldlr; oIMR0092 (mutant): 5’ AATCCATCTTGTTCAATGGCCGATC, oIMR3349 (common): 5’ CCATATGCATCCCCAGTCTT, oIMR3350 (wild-type): 5’ GCGATGGATACACTCACTGC, LysMcre; oIMR3066 (mutant): 5’ CCCAGAAATGCCAGATTACG, oIMR3067 (common): 5’ CTTGGGCTGCCAGAATTTCTC, oIMR3068 (wild-type): 5’ TTACAGTCGGCCAGGCTGAC.

#### Diet induced atherosclerosis

Eight-week old WT^Ldlr-KO^ and M-S196A^Ldlr-KO^ male mice were fed ad libitum a Western diet (WD) (20% Fat, 0.15% Cholesterol; #5342 AIN-7A, Test Diet Limited, UK) for 12 weeks. Mice were fasted overnight prior to sacrifice.

### Metabolic tests

Blood was sampled from saphenous vein as previously described(59). Glucose concentrations were determined in whole blood by a portable meter (Roche Diagnostics, 2 Burgess Hill, UK). Plasma insulin concentrations were determined by enzyme-linked immunoassay (#EZRMI-13K, Merck Millipore). Plasma total cholesterol and triglyceride levels (Wako Diagnostics), as well as NEFAs (Abcam), were determined by colorimetric enzymatic assay kits as per the manufacturer’s recommendations.

### Atherosclerosis quantification

#### *En face* analysis of aorta

Mice were perfusion-fixed with phosphate-buffered paraformaldehyde (4% [wt/vol.], pH 7.2) under terminal anaesthesia. The entire aortic tree was dissected free of fat and other tissue. Aortae were stained with oil red O and mounted onto glass slides before imaging (Leica, DFC310FX) under a dissection microscope (Leica, MZ10F). Lesion area of whole aorta was analysed using Image J.

#### Aortic sinus

Hearts were formalin-fixed, paraffin-embedded and 5 μm aortic sinus sections were stained with stained with hematoxylin and eosin (H&E). Stained sections were scanned with NanoZoomer Digital slide scanner (Hamamatsu). Percentage atherosclerotic lesion were determined using Image J by averaging 3 sections from each mouse with 30-50 μm intervals between sections(59).

### Macrophage content

Immunohistochemistry staining was performed at the UCL IQPath Laboratory using the Ventana Discovery XT instrument, using the Ventana DAB Map detection Kit (760-124). For pre-treatment Ventana Protease 1 (equivalent to pronase, 760-2018) was used. CD68 primary antibody (AbD Serotec #MCA1957), followed by Rabbit anti Mouse (#E0354 Dako). Stained sections were scanned with NanoZoomer Digital slide scanner (Hamamatsu). Macrophage content was quantified in the plaque using Image J.

#### Plaque complexity

Percent necrotic core was measured in H&E stained aortic roots as acellular area(60) using Image J.

### Cell culture

#### Bone marrow derived macrophage culture

Bone Marrow-derived Macrophages (BMDM) were prepared as in(61) using L929 Conditioned Medium (LCM) as a source of M-CSF for the differentiation of the macrophages. After 6 days of differentiation, LCM-containing medium was removed, cells were washed three times in warm PBS and incubated in DMEM containing low-endotoxin (≤10 EU/mL) 1% FBS and 20 μg/mL gentamycin without any LCM before being treated with DMSO or 1 μmol/L GW3965 (Tocris) for 24 hrs.

#### Isolation of mouse peritoneal macrophages

Peritoneal macrophages were harvested 4 days after i.p. injection of 4% thioglycolate by peritoneal lavage. Macrophages were seeded at 2×10^6^ cells/mL in RPMI and adherent macrophages were washed in PBS and harvested after 2 hours.

#### Jurkat culture

Jurkat cells were cultured in RPMI supplemented with 10% FBS, and passaged every two days to maintain a cell concentration not exceeding 1×10^6^ cells/mL.

### Efferocytosis

Jurkat cells or BMDM were labeled for 1 hr with calcein AM and apoptosis was induced by UV irradiation. Apoptotic cells were added to monolayers of BMDM at a ratio of 1:1. After 30 min of coculture, non-ingested apoptotic cells were removed, and slides fixed in 1% PFA. Images were captured (microscope: Zeiss Axio Vert.A1, camera: Zeiss Axiocam 503 mono) and ingested apoptotic cells quantified as in(62) using Image J.

### Ki67 Staining by flow cytometry

Macrophages were washed with PBS before incubation with dissociation media (PBS + 10μM EDTA + 4mg/mL lidocaine + Pen/Strep) at 37°C for 15 minutes and removal from tissue culture plates by gentle scraping. Cells were stained with a Zombie^™^ Fixable Viability Dye (BioLegend), followed by anti-Mouse F4/80-FITC (Clone BM8, eBioscience^TM^), then fixed and permeabilised using eBioscience^™^ Fixation/Permeabilisation reagents (Invitrogen). Intracellular staining with PE Mouse anti-Ki67 Set (BD Pharmingen^TM^) was performed in eBioScience^™^ Permeabilisation Buffer (Invitrogen) for 30 minutes at 4°C. Samples were acquired on a BD LSR Fortessa^™^ X-20 (BDBioscience) using BD FACSDiva^™^ Software. Data was analysed using FlowJo^®^ v10.4 (Tree Star Inc.)

### RNA extraction and quantification

Total RNA from was extracted with TRIzol Reagent (Invitrogen). Sample concentration and purity was determined using a NanoDrop^™^ 1000 Spectrophotometer and cDNA was synthesized using the qScript cDNA Synthesis Kit (Quanta). Specific genes were amplified and quantified by quantitative Real Time-PCR, using the PerfeCTa SYBR Green FastMix (Quanta) on an MX3000p system (Agilent). Primer sequences are available upon request The relative amount of mRNAs was calculated using the comparative Ct method and normalized to the expression of cyclophylin(63).

### Protein isolation and immunoblotting

Total cellular protein lysates (30μg) were loaded onto a 10% SDS-PAGE gel, electrophoresed and transferred onto a PVDF membrane. The membrane was probed with anti-FoxM1 (G-5: sc-376471, Santa Cruz), anti-Akt and anti-phosphorylated Akt (#8200S, Cell Signaling) and anti-Hsp90 (sc-7947, Santa Cruz) overnight in 2.5% BSA, TBS, followed by incubation with anti-rabbit (PO448, Dako) or anti-mouse (NA931VS, GE Healthcare) horseradish-peroxidase-tagged antibodies. Chemiluminescence (ECL 2 Western Blotting Substrate, Pierce) was used to visualise proteins.

### RNA sequencing and analysis

Total RNA was extracted using TRIzol reagent (Life technologies) and cDNA libraries were prepared using reagents and protocols supplied with the Stranded mRNA-Seq Kit (Kapa Biosystems). Briefly, poly-A tailed RNA was purified using paramagnetic oligo-dT beads from 200 nanograms of total RNA, with a RNA Integrity Number above 7.5 as determined by the Agilent Bioanalyzer (Agilent). The purified RNA was chemically fragmented and cDNA was synthesised using random primers (Kapa Biosystems). Adapter-ligated DNA library was amplified with 12 cycles of PCR and library fragment was estimated using the Agilent TapeStation 2200.Library concentration was determined using the Qubit DNA HS assay (Life Technologies). Libraries were sequenced on an Illumina NextSeq 500, NCS v2.1.2 (Illumina) with a 43bp paired end protocol. Basecalling was done using standard Illumina parameters (RTA 2.4.11). Sequencing and pipeline analysis was performed by UCL Genomics (London, UK). Reads were demulitplexed using Illumina bcl2fastq v2.17 (Illumina) and aligned using STAR v2.5.0b to the mouse GRCm38/mm10 reference sequence. Transcript abundance was estimated using Illumina’s RnaReadCounter tool and differential expression analysis performed with DESeq2, which uses the Benjamin-Hochberg method for multiple testing correction. Pathway enrichment analysis was performed with the Gene Set Enrichment Analysis (GSEA) software’s preranked module(64, 65) or the GSEA module in the WebGestalt (http://www.webgestalt.org) analysis toolkit. Reactome pathway analysis was performed with WebGestalt using the Benjamini-Hochberg method to adjust p values for multiple testing and FDR <0.05. Heatmaps of gene counts were done with Heatmapper Expression tool(66) (http://www1.heatmapper.ca/expression/) and Venn diagrams using a BGE tool (http://bioinformatics.psb.ugent.be/webtools/Venn/).

### ChIP-sequencing

Immortalized bone marrow derived-macrophages (iBMDM) from LXRαβ^-/-^ mice have been described(67). N-terminus 3xFLAG-tagged LXRα or LXRβwere ectopically expressed in LXR-null iBMDM using a pBabe-puro retroviral expression system as described(68). A control LXRαβ^-/-^ iBMDM line was also prepared by transduction with an empty pBabe-puro vector. For genome-wide binding analysis of LXR proteins, FLAG-tagged cells were cultured in DMEM supplemented with 1% FFA-free BSA, 50nM of Zaragozic acid (Squalene Synthase inhibitor; Sigma), and 1 uM of GW3965 LXR agonist or GW2033 LXR antagonist (both kindly provided by Jon Collins, Glaxo SmithKline) for 24 h. Immortalized bone marrow derived-macrophages (12 × 10^6^) were crosslinked with 2mM DSG (disuccinimidyl glutarate) for 30 min and 1% methanol-free ultrapure formaldehyde for 10 min before quenching with 2 M Glycine. Cells were lysed with RIPA buffer and, after chromatin shearing by sonication (Bioruptor Diagenode), incubated overnight with protein G magnetic Dynabeads (Invitrogen) that were previously coupled with 3 μg of either anti-FLAG M2 (Sigma) or anti-H3K27ac (Abcam #ab4729) antibodies according to the manufacturer’s instructions. Immunoprecipitated DNA was purified using Qiagen columns. For high-throughput sequencing, ds DNA was obtained by pooling DNA from 10 independent ChIP (for FLAG-LXR sequencing) or 6 different ChIP (for H3K27ac sequencing). DNA was then used for library preparation and subsequent Illumina HiSeq sequencing by the Centre de Regulació Genomica (CRG, Barcelona, Spain) genomic facility.

#### Statistics

Results are expressed as mean (SEM). Comparisons within groups were made using paired Students t-tests and between groups using unpaired Students t tests or repeated measures ANOVA, as appropriate; where repeated *t-tests* were performed a Bonferroni correction was applied. P≤0.05 considered statistically significant except for RNAseq analysis where P≤0.01 was used.

## AUTHOR CONTRIBUTIONS

M.G. performed most experiments, data analysis and prepared figures. N.B. performed experiments and helped with RNAseq analysis. L.R., K.W., T.T. and A.Y. performed qPCR and analysed data. K.W. performed Ki67 proliferation analysis. B.P. and O.M.P. helped establish mouse colonies. J.V.R and A.C. performed ChIPseq analysis. Z.Y. performed efferocytosis assays. M.G. and I.P.T designed experiments, analysed and interpreted data and wrote the manuscript. I.P.T. conceived the study, secured funding and supervised all aspects of the work.

## ACKNOWLEDGEMENTS

We are grateful to Prof Edward Fisher and Prof Michael Garabedian (NYU School of Medicine) for their insightful discussions and Theresa Leon for providing Jurkat cells. This work was supported by a Medical Research Council New Investigator Grant G0801278 (IPT), British Heart Foundation Project Grant PG/13/10/30000 (IPT), UCL Grand Challenges PhD Studentship (IPT), BHF Studentship FS/12/59/30649 (IPT), and the Royal Free Charity PhD Program in Medicine (IPT), and SPANISH MINECO grants SAF2014-56819-R (AC), SAF2015-71878-REDT (AC).

## REFERENCES

1. Roth GA, et al. (2015) Global and regional patterns in cardiovascular mortality from 1990 to 2013. Circulation 132(17):1667–78.

2. Moore KJ, Sheedy FJ, Fisher E a (2013) Macrophages in atherosclerosis: a dynamic balance. Nat Rev Immunol 13(10):709–21.

3. Lee SD, Tontonoz P (2015) Liver X receptors at the intersection of lipid metabolism and atherogenesis. Atherosclerosis 242(1):29–36.

4. Pascual-García M, Valledor AF (2012) Biological Roles of Liver X Receptors in Immune Cells. Arch Immunol Ther Exp (Warsz) 60(4):235–249.

5. Becares N, Gage MC, Pineda-Torra I (2017) Posttranslational Modifications of Lipid-Activated Nuclear Receptors: Focus on Metabolism. Endocrinology 158(2):213–225.

6. Janowski BA, Willy PJ, Devi TR, Falck JR, Mangelsdorf DJ (1996) An oxysterol signalling pathway mediated by the nuclear receptor LXRα. Nature 383(6602):728–731.

7. Spann NJ, et al. (2012) Regulated accumulation of desmosterol integrates macrophage lipid metabolism and inflammatory responses. Cell 151(1):138–52.

8. Collins JL, et al. (2002) Identification of a nonsteroidal liver X receptor agonist through parallel array synthesis of tertiary amines. J Med Chem 45(10):1963–1966.

9. Hong C, Tontonoz P (2014) Liver X receptors in lipid metabolism: opportunities for drug discovery. Nat Rev Drug Discov 13(6):433–444.

10. Ito A, et al. (2015) LXRs link metabolism to inflammation through Abca1-dependent regulation of membrane composition and TLR signaling. Elife 4:e08009.

11. Feig JE, et al. (2010) LXR promotes the maximal egress of monocyte-derived cells from mouse aortic plaques during atherosclerosis regression. J Clin Invest 120(12):4415–4424.

12. Bischoff ED, et al. (2010) Non-redundant roles for LXR and LXR in atherosclerosis susceptibility in low density lipoprotein receptor knockout mice. J Lipid Res 51(5):900–906.

13. Torra IP, et al. (2008) Phosphorylation of liver X receptor alpha selectively regulates target gene expression in macrophages. Mol Cell Biol 28(8):2626–36.

14. Wu C, et al. (2015) Modulation of macrophage gene expression via LXRα serine 198 phosphorylation. Mol Cell Biol 35(11):2024–2034.

15. Ley K, Pramod AB, Croft M, Ravichandran KS, Ting JP (2016) How Mouse Macrophages Sense What Is Going On. Front Immunol 7:204.

16. Nag A, Chakraborty S Biology of FOXM1 and its Emerging Role in Cancer Therapy. J Proteins proteomics.

17. Hu C, et al. (2014) LXRα-mediated downregulation of FOXM1 suppresses the proliferation of hepatocellular carcinoma cells. Oncogene 33(22):2888–2897.

18. Teboul M, et al. (1995) OR-1, a member of the nuclear receptor superfamily that interacts with the 9-cis-retinoic acid receptor. Proc Natl Acad Sci U S A 92(6):2096–100.

19. Tang J, et al. Inhibiting macrophage proliferation suppresses atherosclerotic plaque inflammation. doi:10.1126/sciadv.1400223.

20. Robbins CS, et al. (2013) Local proliferation dominates lesional macrophage accumulation in atherosclerosis. Nat Med 19(9): 1166–1172.

21. Tang J, et al. (2015) Inhibiting macrophage proliferation suppresses atherosclerotic plaque inflammation. Sci Adv 1(3):e1400223–e1400223.

22. Lhoták Š, et al. (2016) Characterization of Proliferating Lesion-Resident Cells During All Stages of Atherosclerotic Growth. J Am Heart Assoc 5(8). Available at: http://jaha.ahajournals.org/content/5Z8/e003945 [Accessed April 4, 2017].

23. Kojima Y, Weissman IL, Leeper NJ (2017) The Role of Efferocytosis in Atherosclerosis. Circulation 135(5):476–489.

24. Kojima Y, et al. (2016) CD47-blocking antibodies restore phagocytosis and prevent atherosclerosis. Nature 536(7614):86–90.

25. Moore KJ, Sheedy FJ, Fisher EA (2013) Macrophages in atherosclerosis: a dynamic balance. Nat Rev Immunol 13(10):709–721.

26. Jenkins SJ, et al. (2011) Local Macrophage Proliferation, Rather than Recruitment from the Blood, Is a Signature of TH2 Inflammation. Science (80-) 332(6035):1284–1288.

27. Sakai M, et al. (1996) The scavenger receptor serves as a route for internalization of lysophosphatidylcholine in oxidized low density lipoprotein-induced macrophage proliferation. J Biol Chem 271 (44):27346–27352.

28. Flaveny CA, et al. (2015) Broad Anti-tumor Activity of a Small Molecule that Selectively Targets the Warburg Effect and Lipogenesis. Cancer Cell 28:42–56.

29. Solt LA, Kamenecka TM, Burris TP (2012) LXR-mediated inhibition of CD4+ T helper cells. PLoS One 7(9):e46615.

30. Pascual-Garcia M, et al. (2011) Liver X Receptors Inhibit Macrophage Proliferation through Downregulation of Cyclins D1 and B1 and Cyclin-Dependent Kinases 2 and 4. J Immunol 186(8):4656–4667.

31. Bensinger SJ, et al. (2008) LXR Signaling Couples Sterol Metabolism to Proliferation in the Acquired Immune Response. Cell 134(1):97–111.

32. Fukuchi J, Kokontis JM, Hiipakka RA, Chuu C, Liao S (2004) Antiproliferative effect of liver X receptor agonists on LNCaP human prostate cancer cells. Cancer Res 64(21):7686–7689.

33. Vedin L-L, Lewandowski SA, Parini P, Gustafsson J-Å, Steffensen KR (2009) The oxysterol receptor LXR inhibits proliferation of human breast cancer cells. Carcinogenesis 30(4):575–579.

34. Uno S, et al. (2009) Suppression of β-catenin signaling by liver X receptor ligands. Biochem Pharmacol 77(2): 186–195.

35. Wang J zhong, Fang Y, Ji W dong, Xu H (2017) LXR agonists promote the proliferation of neural progenitor cells through MEK-ERK pathway. Biochem Biophys Res Commun 483(1):216–222.

36. Zhang X, et al. (2012) Cholesterol metabolite, 5-cholesten-3β-25-diol-3-sulfate, promotes hepatic proliferation in mice. J Steroid Biochem Mol Biol 132(3-5):262–270.

37. Teh M-T, et al. (2002) FOXM1 is a downstream target of Gli1 in basal cell carcinomas. Cancer Res 62(16):4773–80.

38. Chan DW, et al. (2008) Over-expression of FOXM1 transcription factor is associated with cervical cancer progression and pathogenesis. J Pathol 215(3):245–52.

39. Curtis C, et al. (2012) The genomic and transcriptomic architecture of 2,000 breast tumours reveals novel subgroups. Nature 486(7403):346–52.

40. Wang I-C, et al. (2008) Transgenic expression of the forkhead box M1 transcription factor induces formation of lung tumors. Oncogene 27(30):4137–4149.

41. Wierstra I (2013) The transcription factor FOXM1 (Forkhead box M1): Proliferation-specific expression, transcription factor function, target genes, mouse models, and normal biological roles. Adv Cancer Res 118:97–398.

42. Korver W, Roose J, Wilson A, Clevers H (1997) The Winged-Helix Transcription Factor Trident is Expressed in Actively Dividing Lymphocytes. Immunobiology 198(1-3): 157–161.

43. Korver W, et al. (1997) The HumanTRIDENT/HFH-11/FKHL16Gene: Structure, Localization, and Promoter Characterization. Genomics 46(3):435–442.

44. Ye H, et al. (1997) Hepatocyte nuclear factor 3/fork head homolog 11 is expressed in proliferating epithelial and mesenchymal cells of embryonic and adult tissues. Mol Cell Biol 17(3):1626–41.

45. Down CF, Millour J, Lam EWF, Watson RJ (2012) Binding of FoxM1 to G2/M gene promoters is dependent upon B-Myb. Biochim Biophys Acta - Gene Regul Mech 1819(8):855–862.

46. Weidmann H, et al. (2015) SASH1, a new potential link between smoking and atherosclerosis. Atherosclerosis 242(2):571–579.

47. Tanaka S, et al. (2016) BubR1 Insufficiency Results in Decreased Macrophage Proliferation and Attenuated Atherogenesis in Apolipoprotein E-Deficient Mice. J Am Heart Assoc 5(9):e004081.

48. Hoeksema MA, de Winther MPJ (2016) Epigenetic Regulation of Monocyte and Macrophage Function. Antioxid Redox Signal 25(14):758–774.

49. Thangavel J, et al. (2015) Epigenetic modifiers reduce inflammation and modulate macrophage phenotype during endotoxemia-induced acute lung injury. J Cell Sci 128(16):3094–105.

50. Sun C, et al. (2012) The phenotype and functional alterations of macrophages in mice with hyperglycemia for long term. J Cell Physiol 227(4):1670–1679.

51. Gallagher KA, et al. (2015) Epigenetic changes in bone marrow progenitor cells influence the inflammatory phenotype and alter wound healing in type 2 diabetes. Diabetes 64(4). doi:10.2337/db14-0872.

52. A-Gonzalez N, et al. (2009) Apoptotic cells promote their own clearance and immune tolerance through activation of the nuclear receptor LXR. Immunity 31 (2):245–58.

53. Thorp E, Cui D, Schrijvers DM, Kuriakose G, Tabas I (2008) Mertk receptor mutation reduces efferocytosis efficiency and promotes apoptotic cell accumulation and plaque necrosis in atherosclerotic lesions of apoe-/-mice. Arterioscler Thromb Vasc Biol 28(8):1421–8.

54. Tanaka T, Terada M, Ariyoshi K, Morimoto K (2010) Monocyte chemoattractant protein-1/CC chemokine ligand 2 enhances apoptotic cell removal by macrophages through Rac1 activation. Biochem Biophys Res Commun 399(4):677–682.

55. Bolick DT, et al. (2009) G2A deficiency in mice promotes macrophage activation and atherosclerosis. Circ Res 104(3):318–327.

56. Hanayama R, et al. (2002) Identification of a factor that links apoptotic cells to phagocytes. Nature 417(6885): 182–187.

57. Ait-Oufella H, et al. (2007) Lactadherin deficiency leads to apoptotic cell accumulation and accelerated atherosclerosis in mice. Circulation 115(16):2168–2177.

58. Takeuchi T, et al. (2002) Flp recombinase transgenic mice of C57BL/6 strain for conditional gene targeting. Biochem Biophys Res Commun 293(3):953–957.

59. Gage MC, et al. (2013) Endothelium-specific insulin resistance leads to accelerated atherosclerosis in areas with disturbed flow patterns: a role for reactive oxygen species. Atherosclerosis 230(1):131–9.

60. Feng B, et al. (2003) Niemann-Pick C heterozygosity confers resistance to lesional necrosis and macrophage apoptosis in murine atherosclerosis. Proc Natl Acad Sci 100(18):10423–10428.

61. Pineda-Torra I, Gage M, de Juan A, Pello OM (2015) Isolation, Culture, and Polarization of Murine Bone Marrow-Derived and Peritoneal Macrophages. Methods Mol Biol 1339:101–9.

62. Thorp E, et al. (2011) Shedding of the Mer Tyrosine Kinase Receptor Is Mediated by ADAM17 Protein through a Pathway Involving Reactive Oxygen Species, Protein Kinase C, and p38 Mitogen-activated Protein Kinase (MAPK)*. doi:10.1074/jbc.M111.263020.

63. Pourcet B, et al. (2016) The nuclear receptor LXR modulates interleukin-18 levels in macrophages through multiple mechanisms. Sci Rep 6(April):25481.

64. Mootha VK, et al. (2003) PGC-1α-responsive genes involved in oxidative phosphorylation are coordinately downregulated in human diabetes. Nat Genet 34(3):267–273.

65. Subramanian A, et al. (2005) Gene set enrichment analysis: a knowledge-based approach for interpreting genome-wide expression profiles. Proc Natl Acad Sci U S A 102(43):15545–50.

66. Babicki S, et al. (2016) Heatmapper: web-enabled heat mapping for all. Nucleic Acids Res 44(W1):W147–W153.

67. Ito A, et al. (2015) LXRs link metabolism to inflammation through Abca1-dependent regulation of membrane composition and TLR signaling. Elife 4(JULY 2015). doi:10.7554/eLife.08009.

68. Chen M, Bradley MN, Beaven SW, Tontonoz P (2006) Phosphorylation of the liver X receptors. FEBS Lett 580(20):4835–4841.

